# *In-silico* Structural and Molecular Docking-Based Drug Discovery Against Viral Protein (VP35) of Marburg Virus: A potent Agent of MAVD

**DOI:** 10.1101/2021.02.09.430405

**Authors:** Sameer Quazi, Javed Malik, Arnaud Martino Capuzzo, Kamal Singh Suman, Zeshan Haider

## Abstract

The Marburg virus (MARV) is a highly etiological agent of hemorrhagic fever in humans. MARV has spread across the world, including America, Australia, Europe, and different Asia countries. However, there is no approved vaccine to combat MARV, combined with a high mortality rate, which makes antiviral drugs against MARV urgent. The viral protein (VP35) is a core protein of MARV that involves multiple functions of the infection cycle. This research used an *in-silico* drug design technique to discover the new drug-like small molecules that inhibit VP35 replication. First, several combinations of ∼ 4260 showed that structure-based similarity above 90% was retrieved from an online “PubChem” database. Molecular docking was performed using AutoDock 4.2, and ligands were selected based on docking / S score lower than reference CID_5477931 and RMSD value between 1-2. Finally, about 50 compounds showed greater bonding producing hydrogen, Van der Waals, and polar interactions with VP35. After evaluating their binding energy strength and ADMET analysis, only CID_ 3007938 and CID_11427396 were finalized, which showed the most vital binding energy and a strong inhibitory effect with MARV’s VP35. The higher binding energy, suitable ADMET, and drug similarity parameters suggest that these “CID_ 3007938 and CID_11427396” candidates have incredible latency to inhibit MARV replication; hence, these strengths led to the treatment of MAVD.

## 1. INTRODUCTION

Marburg virus (MARV) is an enveloped virus that is a member of the Filoviridae family. MARV is a non-segmented and single-strand negative RNA of 17kb to 19kb size genome (Bausch et al. 2006; Carroll et al. 2013). MARV has induced intermittent diseases in limited numbers of persons in Africa for ten years following its first discovery in 1968. Two significant events in the Democratic Republic of the Congo demonstrated approximately 83% mortality (Towner et al. 2006; Zhu et al. 2017). Thus, MARV is caused by the disease commonly known as homographic fever in humans and animals (Mehedi et al. 2011). No commercially authorized vaccinations or therapeutics are presently approved to manage MARV infections, and work on MARV is therefore desperately required (Anthony and Bradfute 2015).

MARV contained seven different genes across the entire genome. Each gene contained the open read frame (ORF) compatible with a wide range of 68-648 nucleotide lengths at the flanking ends (Wang et al. 2001). The five structural proteins, including nucleoprotein N.P., the viral proteins (V.P.) 35 and 40, the glycoprotein, and the RNA-dependent RNA polymerase (L), are playing an essential role in the infectivity of MARV (Biacchesi et al. 2003). N.P. performs a pivotal function in the growth and development of virions in MARV. N.P. combines with some other viral proteins, particularly VP24, VP40, VP35, and VP30, as an essential component of the virus assembly machinery to coordinate the replication process (Bamberg et al. 2005; Becker et al. 1998). So, it arranges as the scaffold of nucleocapsid development into a helical tubular structure. Sequence homology reveals that N.P. Includes a preserved N-terminal region appropriate for assembling self-assembly and single-stranded RNA (ssRNA) and a mostly unorganized C-terminal region containing a part necessary for the flourishing of virions (Dolnik et al. 2010; Kolb et al. 2014).

Further, through multipurpose VP35, which plays an essential function in the synthesis of viral RNA, assembly, and structure of the virus, MARV often counteracts immune response. MARV VP35 communicates with many innate antiviral defense elements, particularly mechanisms that contribute to the IFN formation of the RIG-I (Retinoic acid-inducible gene-I) like receptor (Ramanan et al. 2012). The FGI-103 (2-(2-(5-(aminomethyl)-1-benzofuran-2-yl) vinyl)-1H-benzimidazole-5-carboximidamide) is the small drug-like compound that has previously classified and reported as an effective drug against VP35 of MARV (Warren et al. 2010).

The VP35 is considered a vital target to synthesize the antiviral drug due to the important role of VP35 in the transcription of MARV. The FGI-103 drug was selected to screen small molecules from the PubChem database using structure similarity-based filtration (More than 90% similarity) to find novel compounds. The CADD and Virtual high throughput screening perform a critical function in drug discovery (Lyne 2002). The bioinformatics techniques, including structure-based drug-like compounds screening from online databases, molecular docking, and molecular dynamic simulation, could be utilized to block the P1 active site of VP35. The current research was designed to novel drug-like substances with greater contact, binding energy, and inhibition effect at the P1 site of MARV VP35 by using computational strategies. The final small molecules of drug-like compounds would have more effective and substantial latent to stop the replication of MARV in the host, which could ultimately help develop and design new drugs to cure and target MAVD.

## 2. MATERIALS AND METHODS

### 2.1 Amino acid sequence retrieval and analysis

The amino acid sequences of VP35 protein were retrieved from The National Center for Biotechnology (NCBI) (https://www.ncbi.nlm.nih.gov/) database. NCBI is a significant and leading public biomedical database and contains different tools for analyzing genomic and molecular information in computational biology (Jenuth 2000). Furthermore, the protein’s primary sequence was analyzed using an online bioinformatics-based tool Expasy-Protparam (https://web.expasy.org/protparam/). The Protparam tool was used to analyze the different physical and chemical parameters of protein, including the molecular weight, isoelectric point, atomic composition, estimated half-life, amino acid composition, aliphatic index, and grand average hydrophobicity (Garg et al. 2016).

### 2.2 Structure prediction, Evaluation, and Validation of protein

The sequence of VP35 protein was utilized to identify the template having more significant similarity in the protein sequence. The protein sequence was used with the Basic Local Alignment Search Tool (Blast) in Protein Data Bank (PDB) and selected the best structure with the highest similarity in sequence. The three-dimensional structure (3D) of the VP35 protein was developed by using MODELLER v9.25 software. The MODELLER software is a desktop-based computational tool used to indicate the homology-based 3D structure of the protein. The most favorable and accurate 3D model was selected based on the DOPE score (Eswar et al. 2006). The quality of the 3D structure of VP35 was assessed and validated by using an online freely available PROCHECK tool (https://servicesn.mbi.ucla.edu/PROCHECK/). The PROCHECK software highlights the stereochemistry of protein (Laskowski et al. 1993).

### 2.3 Formation of Coordinate files

The 3D structure of the VP35 protein was modified by using bioinformatics software’s Discovery Studio Visualizer and AutoDock4.2. The structure was optimized by removing the water molecules from the VP35, hydrogen and polar hydrogen atoms, the addition of Kollman charges and fixed the receptor atoms. Finally, VP35 structure was saved in “Pdbqt” format file (Haider et al. 2020; Quazi et al. 2021).

### 2.4 Selection of ligand and database virtual screening

The “. SDF” file of the FGI-103 antiviral drug was downloaded from the PubChem database. The active site (P1) of VP35 was categorized by using an online DoGSite (https://proteins.plus/help/dogsite). The P1 “ALA210,14, LYS211, LEU-215, PHE-218, ILE-230, GLN-233, VAL-234, SER-236, LYS-237, VAL-280, PRO-282, ILE-284, and CYS-315”, of selected for the molecular docking with FGI-103 antiviral drug-using AutoDock 4.2 software. After that, FGI-103 was set and screen other drug-like compounds from PubChem databases. The Pfizer law was used to evaluate the drug-like properties of each compound. The different parameters of Lipinski’s rule like M.W. < 500 Da, LogP < 5, HBD < 10, and HBA < 5, were used to screen the drug-like small molecules (Chen et al. 2020; Lipinski 2004). Selected compounds were nominated for further analysis. Every selected drug-like compound’s energy minimization was completed using AutoDock 4.2 software and saved files into a “pdbqt” file separately for further molecular docking.

### 2.5 Molecular docking

The finally selected drug-like compounds were docked with the P1 site of VP35 of MARV using a desktop AutoDock 4.2 software. The molecular docking was carried out on a computer system that installed a window 10 with an 86x operating system. The applications, including AutoDock 4.2 and MGL 1.5.4 using Python 2.7, were used for these experiments (France, Scotti, and Scotti 2019). The protein-ligand interaction investigation was accomplished utilizing Discovery Studio Visualizer and PyMol software’s, respectively (D Studio and 2008; Inwood et al. 2009). For molecular docking, the receptor and ligand were used after their energy minimization, and both the structure were saved in “. pdbqt” files. The grid chart of all kinds of atom’s energy was generated using AutoGrid algorithm of AutoDock 4.2. A grid box was drawn based on ap1 site for ligand in every dock for VP35 MARV utilizing a grid chart of 50 × 50 × 50 points, 70 × 70 × 70 grid spacing points, and 0.38 Å and 0.44 Å, individually. The docking was completed by selecting the parameters that our previous study described (Haider et al. 2020; Quazi et al. 2021). The “S” value is showed the docking score between the respective receptor and ligand. The more negative “S” indicates the strong binding affinity of the ligand with the receptor. The RMSD (Root Mean Square Deviation) value is utilized the docked conformations between the ligand and receptors. All the docking score and RMSD values of each ligand were calculated using the default scoring parameter in the AutoDock 4.2 (Haider et al. 2020). The “S” score and binding energy of finalized compounds were compared with the values of FGI-103. The small molecule with binding energy like or greater than the FGI-103 was selected. The finally selected compounds were considered for further analysis.

### 2.6 ADMET Profiling

The ADMET properties of finally selected drug-like compounds were checked to utilize an available admetSAR (<u>Immd.ecust.edu.cn/admetsar2</u>) tool. This admetSAR expects multiple toxic effects, including mutagenicity, annoyance behaviour, and competitiveness. The ADMET profile’s drug-like properties help pick healthy human antiviral medicines (Fatima et al. 2020).

## 3. RESULTS

### 3.1 Amino acid sequence retrieval and analysis

The VP35 primary sequence of 329 Amino Acids (A.A.) was obtained from the NCBI database. The stability of the protein structure has relied on the three-dimensional conformation of the protein. The protein sequence of the target protein was developed based on physical and chemical properties. The physicochemical properties estimated using Expasy-Protparam showed that the Molecular weight (W.W.) of protein 36201.85, isoelectric point (pI), 8.84, and Grand average hydropathicity −0.193. All the physical and chemical properties of VP35 protein are shown in Table 1.

**Table 1.**
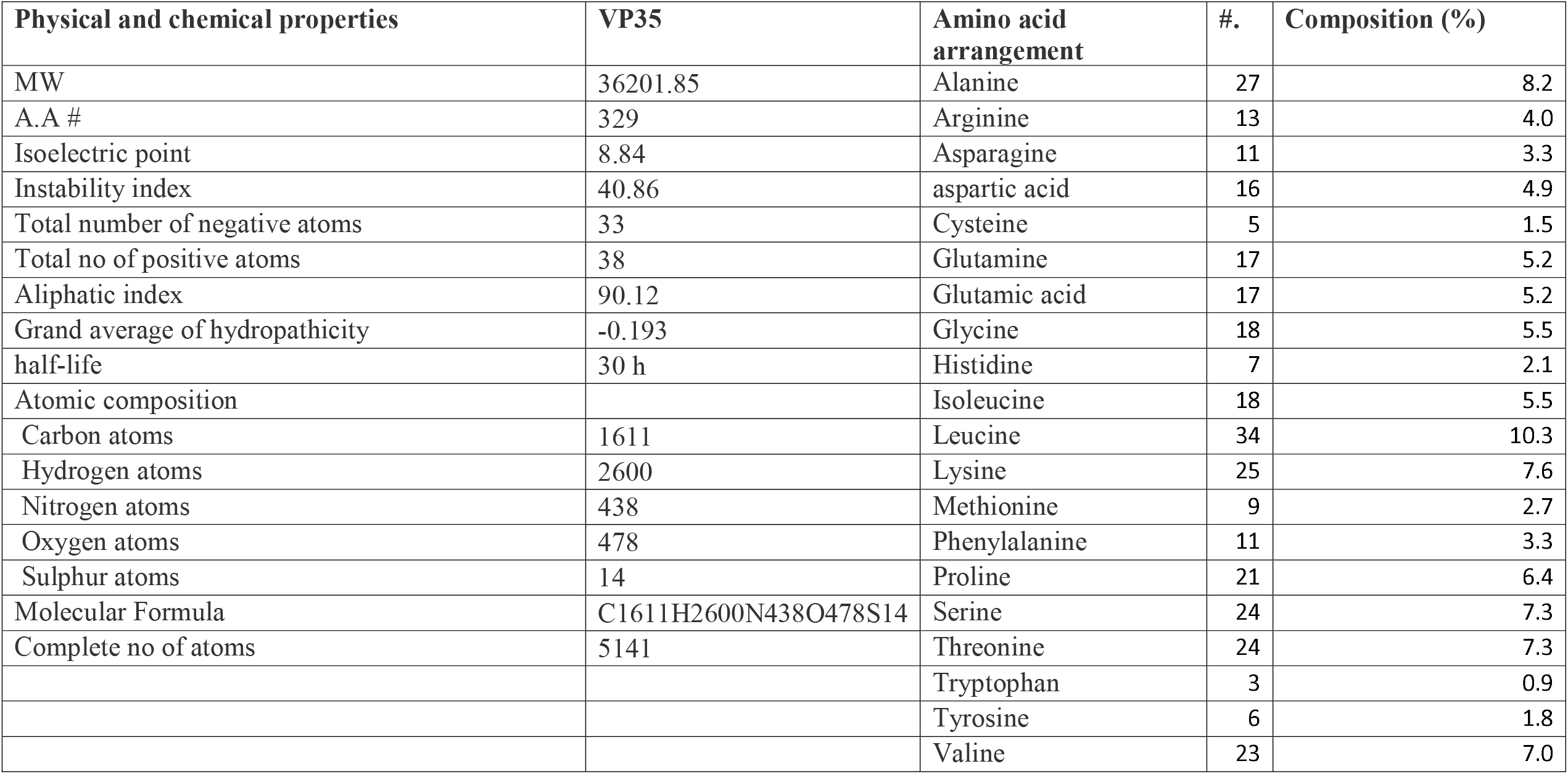
Physio-chemical characteristics of VP35 MARV protein

### 3.2 Structure prediction, Evaluation, and Validation of protein

The 3D structure of the VP35 protein was predicted homology-based. The template “c4gh9A” showed 85% sequence similarity downloaded from PDB. The homology modelling was done by using MODELLER desktop software. The finest 3D structure of VP35 out of ten structures was chosen based on a DOPE and G.A. score (−33763.4352 and 336) (Figure 1). The geomaterial analysis predicted 3D VP35 was performed using PRICHECK tool that showed most of the A.A. approximately 319 A. An out 329 were situated in the protein’s favorable region that made the 97.1% out of 100%. Moreover, the 3D structure of VP35 was considered more reliable, efficient, and stable for further study (Figure 2).

**Figure 1.**
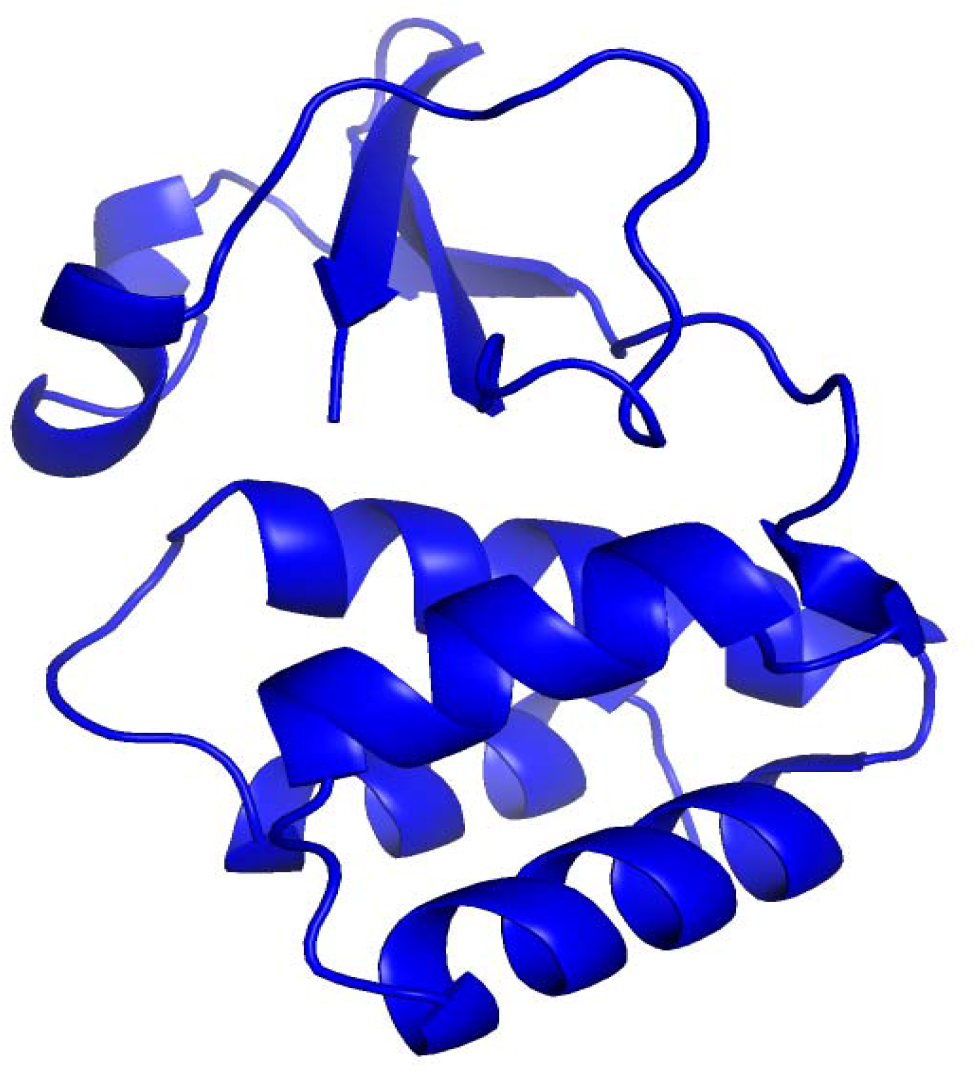
The 3D structure of VP35 by using template “c4gh9A” predicted by PyMol.

**Figure 2.**
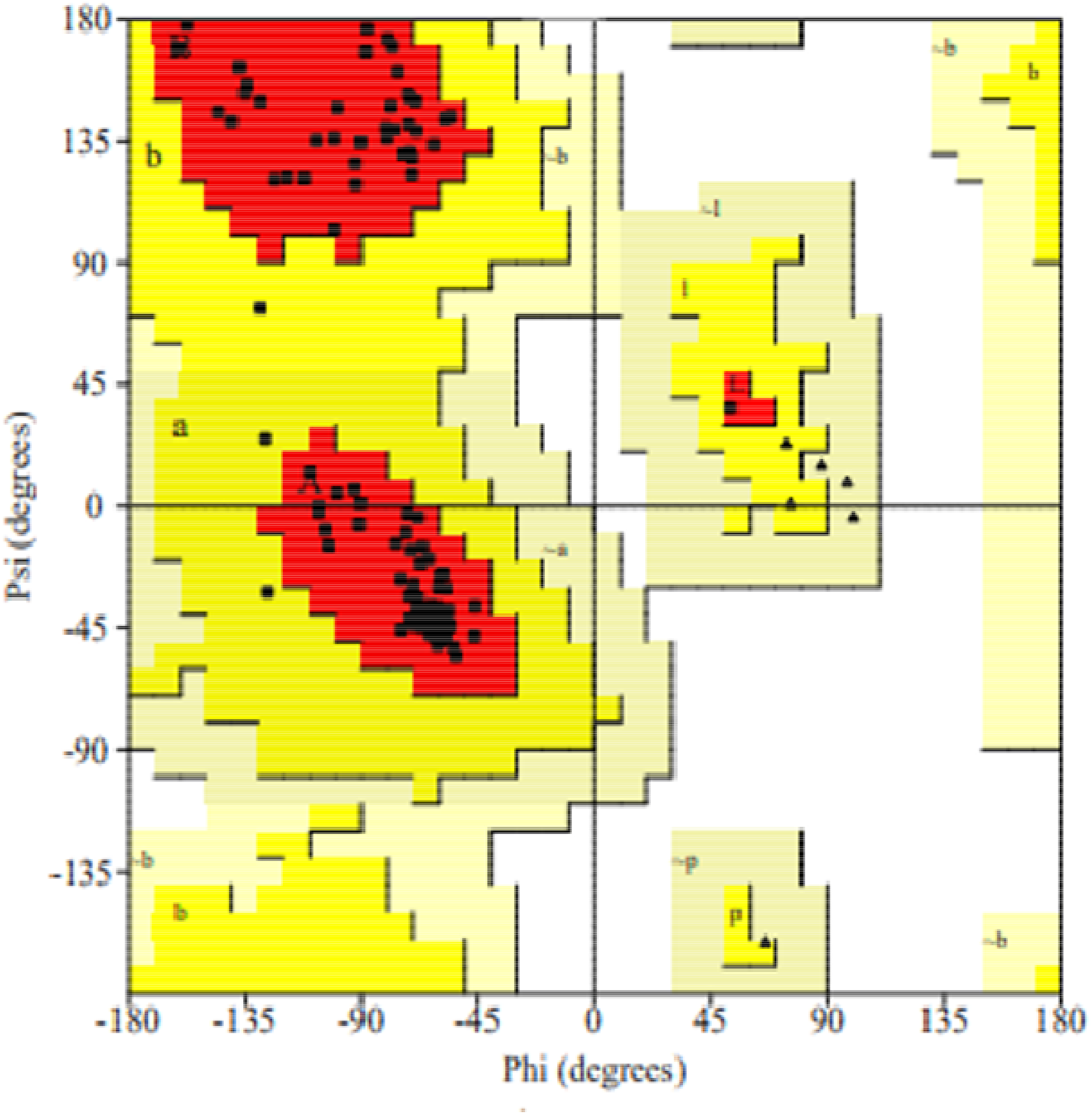
Assessment of Ramachandran’s plot of MARV VP35 shows that 97.1% A. A is present in favorable regions, while about 2.5% A.A, an extant in allowed areas, and 0.4% A.A, exists in an outlier region.

### 3.3 Selection of ligand and database virtual screening

The VP35 protein was docked through an antiviral medication named FGI-103. The results indicated that FGI-103 and the MARV were recognized as being correlated with one another. This analysis showed that the FGI-103 formed a complex with VP35 by “S” score of - 13.46, RMSD of 1.53, and binding energy of - 16.70 (Table 2). The interaction analysis showed that GLN233 formed a strong contact by hydrogen bonding, and GLN230 made a strong connection through polar interaction with FGI-103. While LYS241, VAL279, ILE230, 238 are involved in Van der Waals interactions (Figure 3). In our research, compounds with >90 % structural similarity to FGI-103 were chosen through virtual screening from the broad online PubChem database. Out of the 4260 compounds, Pfizer did cross-validation and applied the law of five to all the combinations in the sample. The 50 out of 4260 drugs like small molecules were placed into another database for docking with the VP35 protein after the most feasible energy minimization.

**Table 2:** The predicted favorable docking results and Pfizer’s properties of finalized drugs compound against VP35.

| PubChem # | Name | Docking score | RMSD | Binding energy kcal/mol | Pfizer's properties |
| --- | --- | --- | --- | --- | --- |
| **CID_5477931** | 1H-Benzimidazole-6-carboximidamide, 2-(2-(5-(aminoiminomethyl)-2-benzofuranyl) ethenyl) | - 13.46 | 1.53 | - 16.70 | MW=347.402, LogP= -1.173, HBD= 6, HBA =0, and tPSA=146.290 |
| **CID_ 3007938** | 2-[4-[5-(4-aminophenyl)-2-furyl] phenyl]-N-isopropyl-3H-benzimidazole-5-carboxamidine<br>1H-Benzimidazole-6-carboximidamide, 2-[4-[5-(4-aminophenyl)-2-furanyl] phenyl]-N-(1-methylethyl)- | -16.33 | 1.85 | -19.45 | MW= 435.53, LogP= 5.85, HBD =3, HBA =2, tPSA=106.22 |
| **CID_11427396** | 2-Dibenzofuran-2-yl-1H-benzoimidazole-5-carboxylic acid amide | -17.78 | 1.54 | -21.23 | MW=329.36, LogP=4.32, HBD =3, HBA=1, tPSA=80.32 |

\*HBD= Hydrogen Bond Donor, HBA =Hydrogen bond acceptor

**Figure 3.**
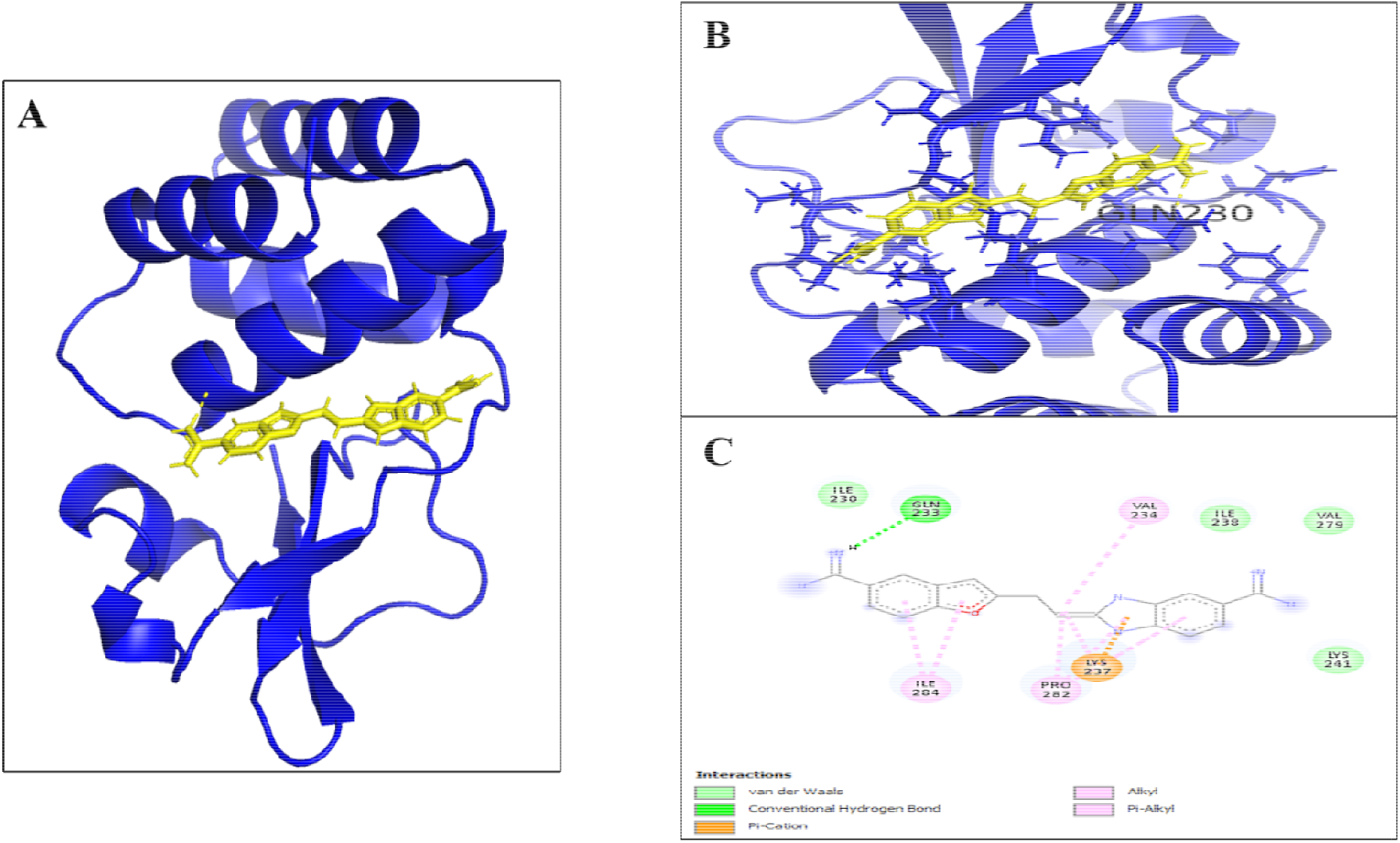
(**A**) 3D structural description of VP35 MARV (Showing in blue colour) formed a complex with FGI-103 (Showing in yellow colour) (**B**) 3D ligand complex between the “ FGI-103” and P1 site of VP35 MARV (**C**) 2D ligand complex between the “ FGI-103” and P1 of VP35 MARV.

## 3.1 Molecular docking

The molecular docking rules play an important role in creating modern drugs against various lethal illnesses (Ursic-Bedoya et al. 2014). All of the hits that were docked against the P1 position of VP35 MARV by utilizing the Auto Dock. Subsequently, there have been two compounds recorded with a minimum S/docking-score than FGI-103. The successful docked top compounds with lower S-score, and RMSD value was selected for further evaluation. The binding relationship of such two-hit compounds with VP35 was determined using Drug Discovery Studio tools. The best locations were defined in the specified order of preference, constructed on the minimum binding energy in the greatest cluster, no. of hydrogen connections formed with A.A, residues of P1 (Table 2). That was done to ensure that the compounds were attached exactly in the correct binding position. Successful inhibitors have shown an important correlation with the P1 site of MARV VP35 (Figure 4).

**Figure 4.**
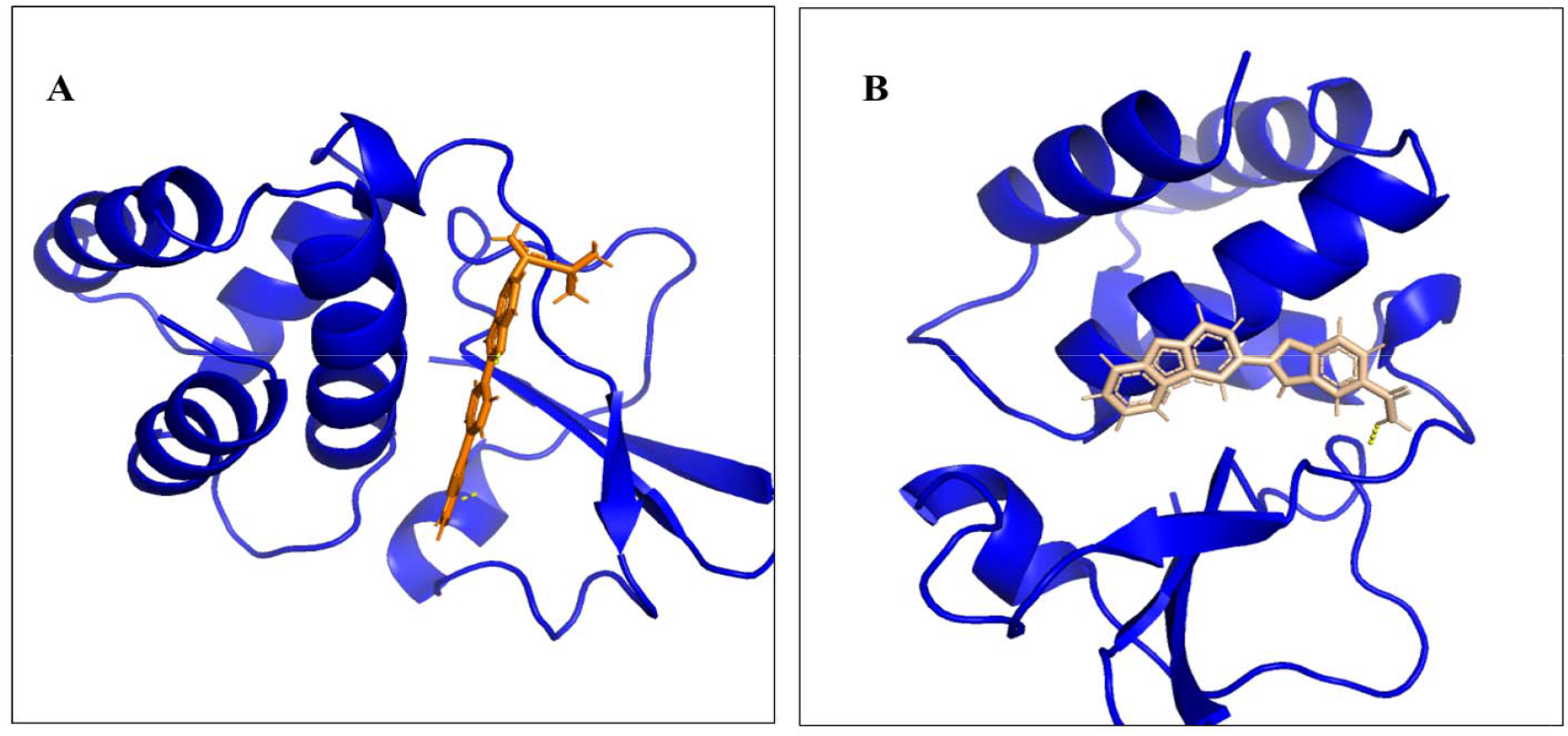
(**A**) Representation of 3D complex of VP35 (Showing in blue colour) interacted with novel inhibitor CID_ 3007938 (Showing in orange colour) (**B**) Representation of 3D complex of VP35 (Showing in blue colour) connected with novel inhibitor CID_11427396 (Showing in tints white colour).

### 3.2 ADMET profiling

The Molinspiration server was used to crisscross the drug-like parameters of suggested small molecules against MARV VP35. The selected compounds showed a zero violation against the Pfizer’s law of five and recognized the properties of drug including M.W., HBD, HBA, LogP and tPSA (Table 2). Further, to evaluate the properties of drugs safety in the living organism. The term ADMET is the abbreviation of Absorption, Digestion, Metabolism, Excretion and Toxicity. The ADMET analysis was performed by using AdmetSAR server. The ADME analysis of all the finally selected compounds showed zero violation against the use in a living organism (Table 3).

**Table 3:** ADME analysis of finalized drugs compounds against VP35.

| PubChem # | Blood-Brain Barrier | Human Intestinal Absorption | CaCO2 permeability | p-glycoprotein inhibitor | Renal Organic Cation Transporter |
| --- | --- | --- | --- | --- | --- |
| **Absorption** |  |  |  |  |  |
| CID_5477931 | Positive (+) | Positive (+) | Negative (-) | Non-Inhibitor | Non-Inhibitor |
| CID_3007938 | Positive (+) | Positive (+) | Negative (-) | Inhibitor | Non-Inhibitor |
| CID_11427396 | Positive (+) | Positive (+) | Negative (-) | Non-Inhibitor | Non-Inhibitor |
| PubChem # | CYP450 1A2 | CYP450 2C9 | CYP450 2D6 | CYP2450 C19 | CYP450 3A4 |
| **Metabolism** |  |  |  |  |  |
| CID_5477931 | Non-Inhibitor | Non-Inhibitor | Non-Inhibitor | Non-Inhibitor | Non-Inhibitor |
| CID_3007938 | Inhibitor | Non-Inhibitor | Non-Inhibitor | Non-Inhibitor | Non-Inhibitor |
| CID_11427396 | Inhibitor | Non-Inhibitor | Non-Inhibitor | Inhibitor | Non-Inhibitor |
| PubChem # | AMES analysis |  | Carcinogenic analysis |  |  |
| **Toxicity** |  |  |  |  |  |
| CID_5477931 | Non-Poisonous |  | Non-dangerous |  |  |
| CID_3007938 | Non-Poisonous |  | Non-dangerous |  |  |
| CID_11427396 | Non-Poisonous |  | Non-dangerous |  |  |

### 3.3 Analysis of receptor-ligand interaction

The S/docking-score tests the contact strength of VP35 against drugs compounds; therefore, the drug-like small molecules are chosen based on the S-score and the binding energy of an outstanding drug compound. The following compounds, CID_ 3007938 and CID_11427396, have a solid binding with the P1 active sites VP35. The receptor-ligand complex of CID_ 3007938 and CID_11427396 with VP35 showed that Hydrogen bonding, Van der Waals, formed a stable complex. The LYS237 made a hydrogen bond along with other amino acids of P1 site formed Van der Waals interaction with CID_ 3007938 ligand (Figure 5 A). while VAL280 formed a hydrogen bond along with other amino acids of P1 site formed Van der Waal interaction with CID_11427396 ligand (Figure 5 B). The 2D and 3D pockets configurations of the particular drug-like small molecules are shown in Figure 5.

**Figure 5.**
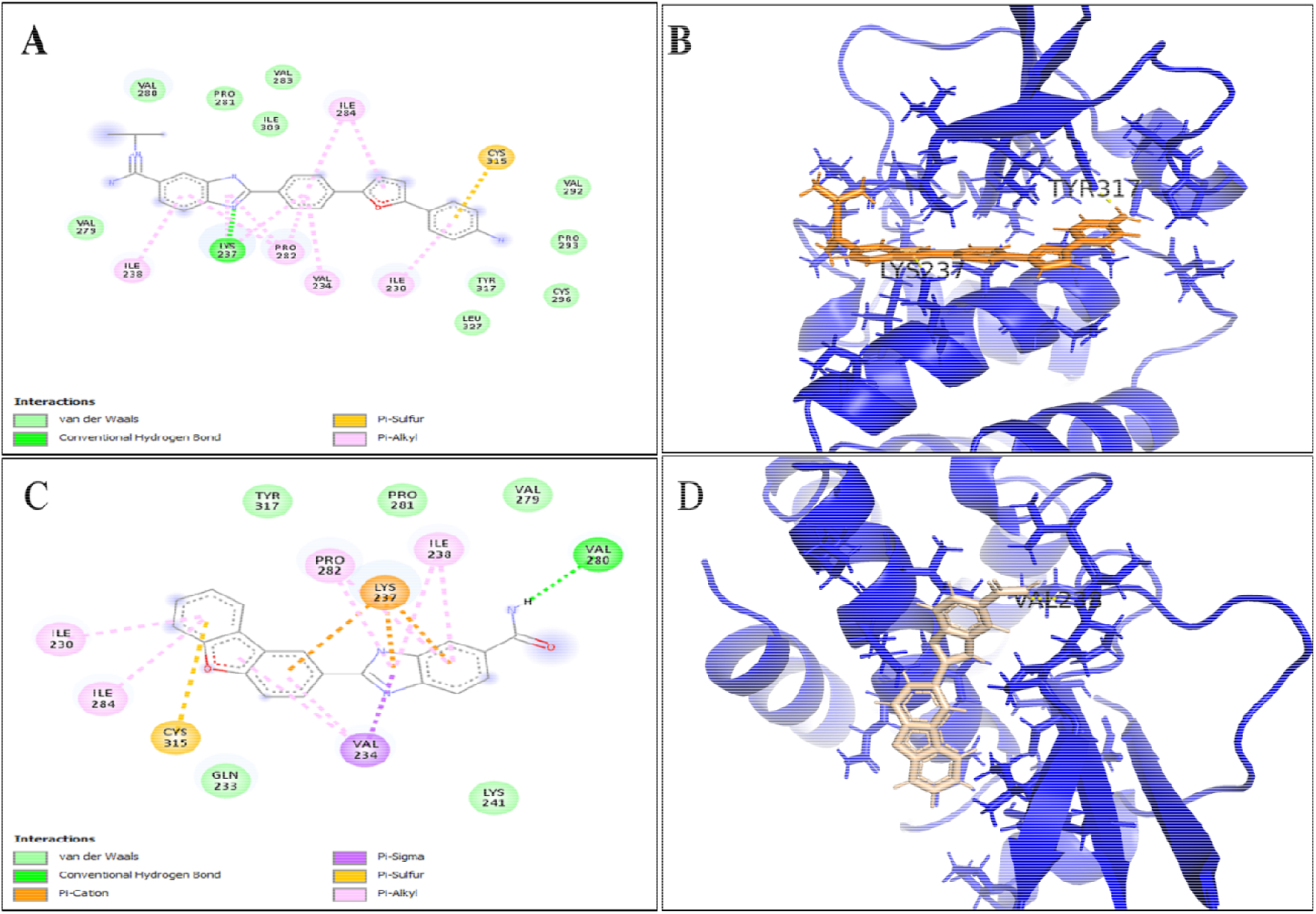
The 2D and 3D representation the analysis of receptor binding interaction (**A**) show the 2D interaction of ligand CID_ 3007938 with VP35 (**B**) represents the 3D pocket of VP35 with Ligand CID_ 3007938 (**C)** shows the 2D interaction of ligand CID_11427396 with VP35. (**D**) represents the 3D pocket of VP35 with Ligand CID_11427396.

## 4. DISCUSSION

Many research-based studies have been conducted to discover effective therapeutic vaccines against MARV. But unfortunately, effective treatment for MARV is not yet available. Nowadays, MARV is considered a global problem and it is still necessary to discover a less expensive and effective antiviral drug against MARV (Brown et al. 2014). Traditional drug development approaches are largely costly and unproductive for solving evolving public health challenges (Velmurugan, Mythily & Rao 2014). Therefore, the most appropriate approaches should be implemented that could easily cope with this adverse circumstance.

In silico drug design strategies are becoming the popular field in the pharmaceutical industries due to fast, less expensive, and time-saving practices in identifying new drugs (Geisbert, Bausch, and Feldmann 2010). MARV’s VP35 viral protein is a promising candidate for vaccine design against MARV infection. Due to the above arguments, current research has proposed small drug-like molecules that caused MARV replication inhibition by binding firmly to the P1 site of VP35 MARV and could be considered pharmacological compounds. The three-small drug-like molecules were also analyzed for ADMET properties with the AdmetSAR server. All the compounds were chosen to have passed the ADMET properties. Blood-brain barrier cells are endothelial cells that function as resistance and prevent the brain from absorbing any medicine. Therefore, blood-brain barrier cells are considered an integral feature in the drug design discipline (Alavijeh et al. 2005; Cheng et al. 2012; Stamatovic, Keep and Andjelkovic 2008). Oral bioavailability is a significant factor for the pharmacological similarity of the active drug compound as a curative agent (Varma et al. 2010). ADMET properties of beneficial drug-like small molecules have strong results for the similarity of effective treatment such as P-glycoprotein substrate (inhibitor / non-inhibitor), blood-brain barrier penetration (positive/negative), human intestinal preparation (positive/negative), renal transporter of organic cations (inhibitor / non-inhibitor) and CaCO2 permeability (positive/negative). Cytochromop450 (CYP) is classified into isoenzymes and has remained active for the catabolism of several chemicals, including hormones, medicines, bile acids, carcinogens, etc. The ADMET research test is useful and efficient for scanning drug compounds and consisted of the following parameters: (1) blood-brain barrier penetration, (2) human intestinal absorption, (3) CaCO2 permeability absorption, (4) non-toxic, (5) non-carcinogenic, and (6) non-inhibitor of the CYP enzyme. These ADMET parameters were significantly exceeded by the two compounds CID 3007938 and CID 11427396 (Table 2).

## 5. CONCLUSION

The current research focus was structure-based virtual screening using the PubChem online database, Pfizer/Lipinski’s analysis, molecular docking, ADMET analysis, and evaluation of the interaction between ligands and the MARV VP35 site P1. The drug-like compounds CID 3007938 and CID 11427396 showed a strong connection with the P1active site of MARV VP35 creating hydrogen bonds, van der Waals and polar interaction. The results suggest that they can hypothetically be applied against MARV as a drug. The compounds mentioned may function as the novel, fundamentally distinct and potentially active pharmaceutical compounds against MARV VP35. The molecule structure of three drug-like compounds is shown in Figure 6. Our in-silico research found that two drugs as small molecules have the potential of a drug that can be guided as therapeutic drugs against MARV by skillfully directing the P1 of VP35 through MARV. Consequently, concerning two small drug-like molecules CID_ 3007938 and CID_11427396, the work we performed requires further investigations and future in vitro and in vivo experiments before a possible verification with the competent authorities.

**Figure 6.**
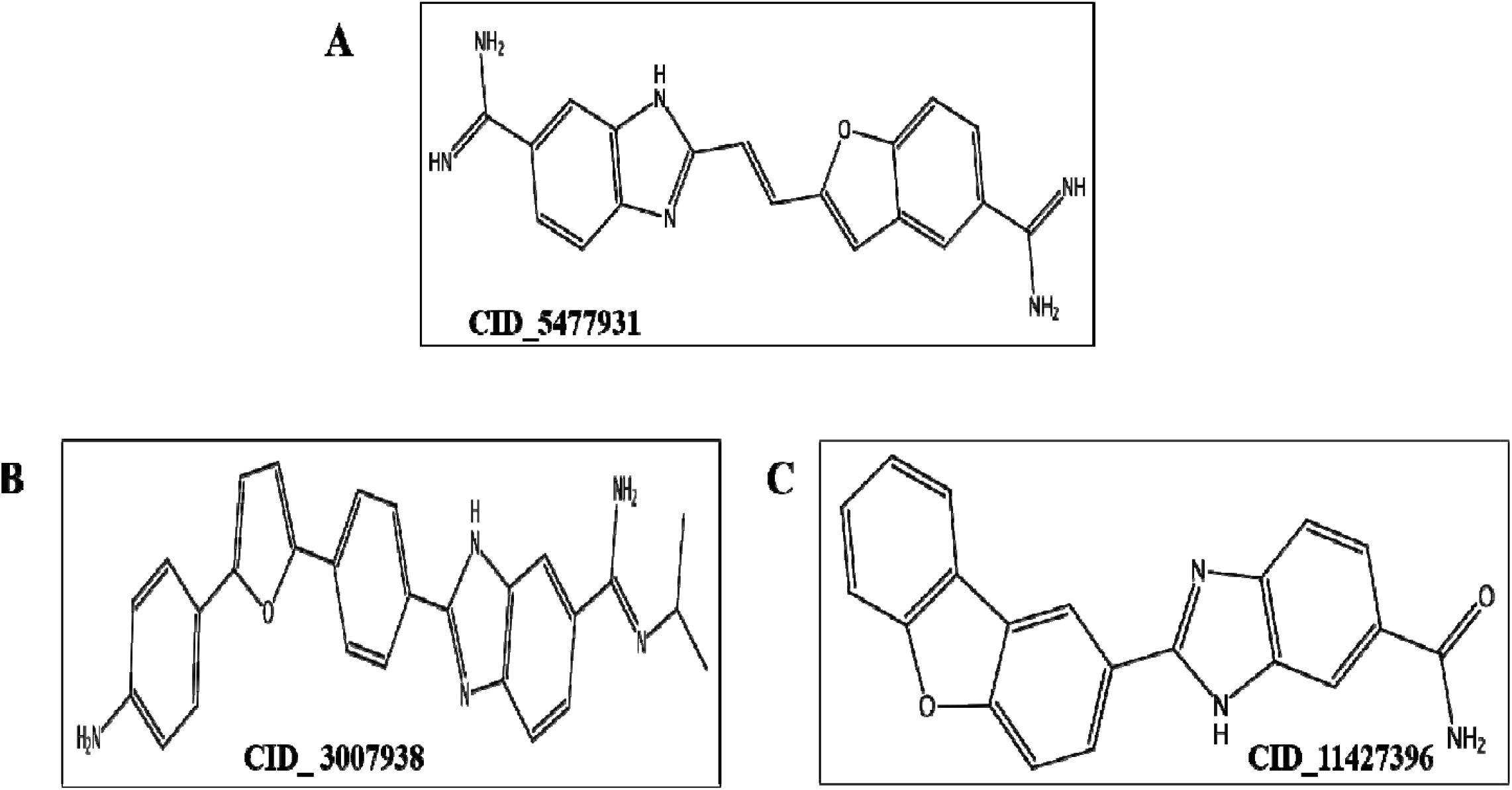
2D molecules structure of selected drug-like compounds **(A)** represents the 2D structure of 1H-Benzimidazole-6-carboximidamide, 2-(2-(5-(aminoiminomethyl)-2-benzofuranyl) ethenyl) drug-like compound **(B)** represents the 2D structure of 2-[4-[5-(4-aminophenyl)-2-furyl] phenyl]-N-isopropyl-3H-benzimidazole-5-carboxamidine 1H-Benzimidazole-6-carboximidamide, 2-[4-[5-(4-aminophenyl)-2-furanyl] phenyl]-N-(1-methylethyl) drug-like compound **(C)** represents the 2D structure of 2-Dibenzofuran-2-yl-1H-benzimidazole-5-carboxylic acid amide drug-like compound

